# Dynamics and mechanisms of ERK activation after different protocols that induce long-term facilitation in *Aplysia*

**DOI:** 10.1101/2022.08.05.502983

**Authors:** Yili Zhang, Rong-Yu Liu, Paul Smolen, Leonard J. Cleary, John H. Byrne

## Abstract

The mitogen-activated protein kinase (MAPK) isoform extracellular signal–regulated kinase (ERK) is a key kinase involved in the induction of long-term synaptic facilitation (LTF) of the *Aplysia* sensorimotor synapse. Therefore, elucidating the dynamics of ERK activation after LTF-inducing protocols is critical for understanding the mechanisms underlying neuronal and synaptic plasticity. ERK activation has rich dynamic features. After a single stimulus, activation peaks 45-minute later and declines rapidly, but after two stimuli spaced 45 min apart, activation persists (Kopec et al. 2015). However, little is known about possible changes in ERK activation for periods beyond 3 h. Given its key role in long-term learning, understanding the dynamics of ERK at 3 h and beyond might provide insights into new protocols that could enhance memory retention. Three different protocols that induce LTF were used to probe the dynamics of ERK activation. The first, termed the Standard protocol, consists of five 5-min pulses of serotonin (5-HT) with regular interstimulus intervals (ISIs) of 20 min. The second, termed the Enhanced protocol, consists of five pulses of 5-HT with irregular ISIs identified with the use of computer simulations. This protocol induces greater and longer-lasting LTF than the Standard protocol. A third protocol, termed the two-pulse protocol, consists of just two 5-min pulses of 5-HT with an ISI of 45 min. Immunofluorescence revealed complex patterns of ERK activation up to 24 h after 5-HT treatment. The Standard and two-pulse protocols led to an immediate increase in active, phosphorylated ERK (pERK), which decayed within 5 h post treatment. A second wave of increased pERK was detected at 18 h post treatment. This late phase was blocked by RpcAMP (an inhibitor of protein kinase A), and by TrkB Fc and TGF-β RII F antagonists. The latter two are chimeras that act via receptor sequestration. These results suggest that complex interactions among kinase pathways and growth factor cascades contribute to the late increase of ERK activity after different LTF-inducing protocols. Interestingly, ERK activity returned to basal levels 24 h after the Standard or two-pulse protocol, but remained elevated 24 h after the Enhanced protocol. This finding may help explain, in part, why the Enhanced protocol is superior to the Standard protocol in inducing long-lasting LTF.

## Introduction

Extensive research has delineated ways in which protein kinase A (PKA) and MAPK pathways contribute to synaptic plasticity that is essential for the formation of long-term memory (LTM) (Abel et al. 1997; Adams and Sweatt 2002; Byrne and Hawkins, 2015; Cleary et al, 1998; English and Sweatt, 1997; Frost et al. 1985; Impey et al. 1998; Kandel 2001; Martin et al. 1997; Matsushita et al. 2001; Mozzachiodi and Byrne 2010; Patterson et al. 2001; Roberson and Sweat 1996; Sharma and Carew, 2004; Vázquez et al. 2000). Our previous studies have combined computational and empirical approaches to investigate how PKA and MAPK cascades contribute to the induction phase of LTF at *Aplysia* sensorimotor synapses (Zhang et al. 2011, 2017, 2021). These studies together with empirical findings from the studies of our and other labs illustrate a high degree of complexity in the dynamics of these kinase pathways and regulation of transcription factors by kinases, including biphasic regulation of kinases and multiple feedback loops, and reveal critical roles played by neurotrophin (NT) and transforming growth factor β (TGF-β) in sustained ERK activation for at least 1 h after 5-HT treatment (Bartsch et al. 1995, 1998; Chin et al. 1999, 2006; Fioravante et al. 2006; Guan et al. 2002; 2003; Jin et al. 2019; Kassabov et al. 2013; Kopec et al. 2015; Liu et al. 1997, 2008, 2011, 2020; Martin et al. 1997; Michael et al. 1998; Mirisis et al. 2020; Müller and Carew, 1998; Ohta et al. 2008; Ormond et al. 2004; Pettigrew et al. 2005; Purcell et al. 2003; Sharma et al. 2003; 2004; Shobe et al. 2016; Song et al. 2007; Zhang et al. 1997, 2017, 2021).

However, most empirical studies have focused on the dynamics of ERK activation within a few hours after 5-HT. Little is known about long-term activation of ERK and how it is regulated. The present study examined these issues using three protocols that induce LTF: the Standard protocol (5 pulses of 5-HT with regular ISIs of 20 min) commonly used to induce LTF for decades (Montarolo et al. 1986; Zhang et al. 2011); the Enhanced protocol (5 pulses of 5-HT with computationally designed irregular ISIs: 10-10-5-30 min) that induced persistent (stronger and longer-lasting) LTF (Zhang et al. 2011); and the 2-pulse protocol with an ISI of 45 min inspired by 2-trial protocols that produce LTM (Kopec et al. 2015; Phillips et al. 2007, 2013). These protocols have substantial differences in the total amount of 5-HT application and duration of treatment.

Here we used immunofluorescence analysis to characterize the dynamics and mechanisms of activation of ERK up to 24 h after the three protocols. Differential active, phosphorylated ERK (pERK) dynamics observed during the late consolidation phase after these protocols may provide further insights into the mechanism underlying the advantage of the Enhanced protocol in producing prolonged LTF, In general, the results emphasize the importance of elucidating the dynamics of kinase cascades for understanding neuronal and synaptic plasticity.

## MATERIALS AND METHODS

### Empirical Methods

#### Neuronal cultures

All experiments used primary cultures of identified sensory neurons (SNs) from *Aplysia californica* (NIH *Aplysia* resource facility, University of Miami, Miami, FL). *Aplysia* are hermaphrodites. Animals were maintained in circulating artificial seawater at 15°C. SNs were isolated from the ventral-caudal cluster of the pleural ganglion from 60–100 gm *Aplysia* according to conventional procedures (Liu et al. 2020; Zhang et al. 2017, 2021). Each dish of SN cultures was plated with 5-10 SNs. SNs were allowed to grow for 5-6 days at 18°C before experiments begun, and the growth medium was replaced at least 2 h prior to treatments with a solution of 50% L15 and 50% artificial seawater (ASW; 450 mM NaCl, 10 mM KCl, 11 mM CaCl_2_, 29 mM MgCl_2_, 10 mM HEPES at pH 7.6).

#### Immunofluorescence analysis

Immunofluorescence procedures for SNs followed those of Zhang et al. (2017). Briefly, after 5-HT treatment, cells were fixed at specific time points in a solution of 4% paraformaldehyde in PBS containing 20% sucrose. Fixed cells were washed by PBS, and then blocked for 30 min at room temperature in a solution of Superblock buffer (Pierce) mixed with 0.2% Triton X-100 and 3% normal goat serum. Cells were subsequently incubated overnight at 4° C with primary anti-phosphorylated ERK antibody (anti-pERK, Cell Signaling, Cat # 4370, RRID: AB_2315112, 1:400). After primary antibody incubation and PBS wash, secondary antibody (goat anti-rabbit secondary antibody conjugated to Rhodamine Red, Jackson ImmunoResearch Lab, Catalog#: 111-295-144, RRID: AB_2338028, 1:200) was applied for 1 h at room temperature. Cells were then mounted using Mowiol 4-88 (SigmaAldrich). The images of cells were obtained with a Zeiss LSM800 confocal microscope using a 63x oil-immersion lens. A z-series of optical sections through the cell body (0.5 μm increments) was taken. The section through the middle of the nucleus was used for quantification of mean fluorescence intensity of the whole cell, with ImageJ-win64 software (NIH). All the cells on each coverslip were analyzed and averaged. The number of samples (n) reported in Results indicates numbers of dishes assessed.

#### Experimental design

For each of the three protocols, a single pulse of 5-HT consisted of a 5-min application of 50 μM 5-HT (Sigma) to SNs. At the end of the 5-min pule, chamber bath was washed with 10 ml of L15/ASW. Dishes of SNs cultured from the same animals were paired for all the 5-HT treatments. One dish received a solution consisting of 50% isotonic L15 and 50% artificial seawater (L15-ASW) as vehicle control (Veh). The other received the same solution with the addition of 5-HT. The experimenter was blind to the identity of treatments given to each pair of dishes.

In the experiments to measure the time course of pERK after 5-HT protocols, each paired dish was either fixed for immunofluorescence immediately after 5-HT, or incubated in L15/ASW after wash off of 5-HT until fixation at specified times (Fig. 2). The remaining dish served as a time-matched Veh control. For each pair of dishes measured at the same time point, the averaged level of pERK from the dish receiving 5-HT was compared to the averaged pERK from the Veh control.

Application of all the inhibitors began 1 h prior to the fixation for immunofluorescence, to ensure that inhibitors were given sufficient time to penetrate the cells and block the activities of kinases. To examine the effects of PKA activity on pERK, 10 μM cAMP inhibitor Rp-cAMP (Calbiochem) was applied to SN cultures 1 h prior to the fixation. Previously, 10 μM cAMP inhibitor Rp-cAMP inhibited the increase of pERK ~45 min post-onset of 5-min treatment with 5-HT without affecting basal activity (Zhang et al. 2021). Four dishes of SNs from the same animals were used for each experiment. Each dish was given a different treatment, either: 1) 50 μM 5-HT alone; 2) 10 μM Rp-cAMP alone; 3) 5-HT + Rp-cAMP; or 4) Veh alone.

To examine the effects of TrkB on pERK, 10 μg/ml of a TrkB antagonist, TrkB Fc chimera (acts via receptor sequestration)(TrkB Fc) (R&D Systems), was applied to SN cultures 1 h prior to fixation. TrkB Fc specifically binds to and neutralizes TrkB ligands, preventing ligand-mediated signaling. Previously, at this concentration, TrkB Fc inhibited the increase of pERK at 1 h after the two-pulse protocol in isolated *Aplysia* SNs without affecting basal activity (Zhang et al. 2021). Four dishes of SNs from the same animals were used for each experiment. Each dish was given a different treatment, either: 1) 5-HT alone; 2) TrkB Fc alone; 3) 5-HT + TrkB Fc; or 4) Veh alone.

To examine the effects of TGF-β on pERK, 5 μg/ml of TGF-β RII Fc chimera (TGF-β RII Fc) (R&D Systems), was applied to SN cultures 1 h prior to fixation to inhibit TGF-β signaling. At this concentration, TGF-β RII Fc inhibited the increase of pERK at 1 h after two pulses of 5-HT (Kopec et al. 2015). Four dishes of SNs from the same animals were used for each experiment. Each dish was given a different treatment, either: 1) 5-HT alone; 2) TGF-β RII Fc alone; 3) 5-HT + TGF-β RII Fc; or 4) Veh alone. The number of samples (n) reported in Results indicates the number of animals.

#### Electrophysiology

Excitatory postsynaptic potentials (EPSPs) were recorded from motor neurons (MNs) from the SN-MN co-cultures following established procedures (Liu et al. 2008, 2011, 2013; Zhang et al. 2012; Zhou et al. 2015). Briefly, the EPSP was evoked in the MN by stimulating the SNs with a brief depolarizing stimulus using an extracellular electrode filled with L15:ASW. Intracellular recordings from MNs were made with 10-20 MΩ sharp electrodes filled with 3 M potassium acetate connected to an Axoclamp 2-B amplifier (Molecular Devices). Data acquisition and analyses of resting potential, input resistance, and EPSP amplitude were performed with pCLAMP 8 software (Molecular Devices). Before measurement of EPSPs, MNs were held at −90 mV by passing constant current. Cultures were excluded from further use if pre-treatment measurements of EPSP amplitudes were less than 10 mV, larger than 35 mV, or sufficiently large to trigger an action potential. MNs with resting potentials more positive than – 40 mV or input resistances less than 10 MΩ were also excluded. These measurements were repeated at 24 h after treatment. The number of samples (n) reported in Results indicates the number of co-cultures.

#### Statistical analyses

At least five animals were used in each experiment. SigmaPlot version 11 (Systat Software) was used for statistical analyses. Before applying statistical tests, Shapiro-Wilk Normality and Equal Variance tests were performed. In the experiments to compare EPSPs between Veh and 5-HT treatment groups, Student t-tests were used (Fig. 1). In the experiments to compare pERK between paired Veh and 5-HT treatment groups at different times, a paired t-test with Bonferroni corrections was used for comparison between paired groups if data passed normality and equal variance tests at all time points. Otherwise, a Wilcoxon Signed Rank Test (WSRT) with Bonferroni corrections was used. In Fig. 2A, because data for the normality variance test failed at one time point, the WSRT with Bonferroni corrections was used for comparison of pERK immunoreactivity between paired Veh and 5-HT treatment groups at all time points Adjusted p values after Bonferroni corrections were used to represent statistical significance.

**Figure 1.**
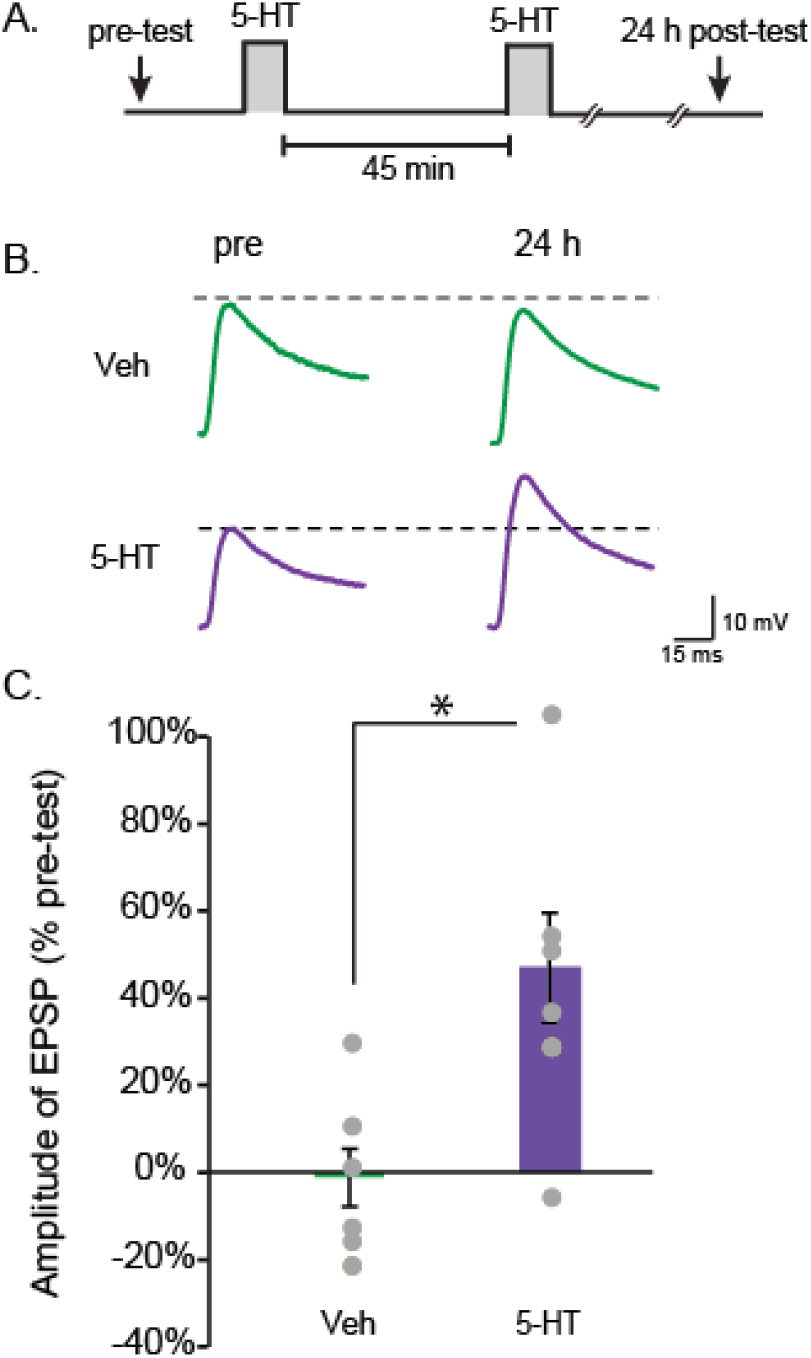
Induction of LTF by the two-pulse protocol. (**A**), Protocol for two pulse treatment with 5-HT (50 μM). (**B**), Representative EPSPs recorded from MNs in sensorimotor co-cultures before (Pre) and 24 h after treatment. (**C)**, Summary data. Two-pulse treatment with 5-HT induced significant increases in the amplitude of EPSPs. Bar height represents the mean, small bars represent standard error of the mean (SEM), and significant differences are indicated by * for p < 0.05. Gray circles indicate the results of individual experiments.

**Figure 2.**
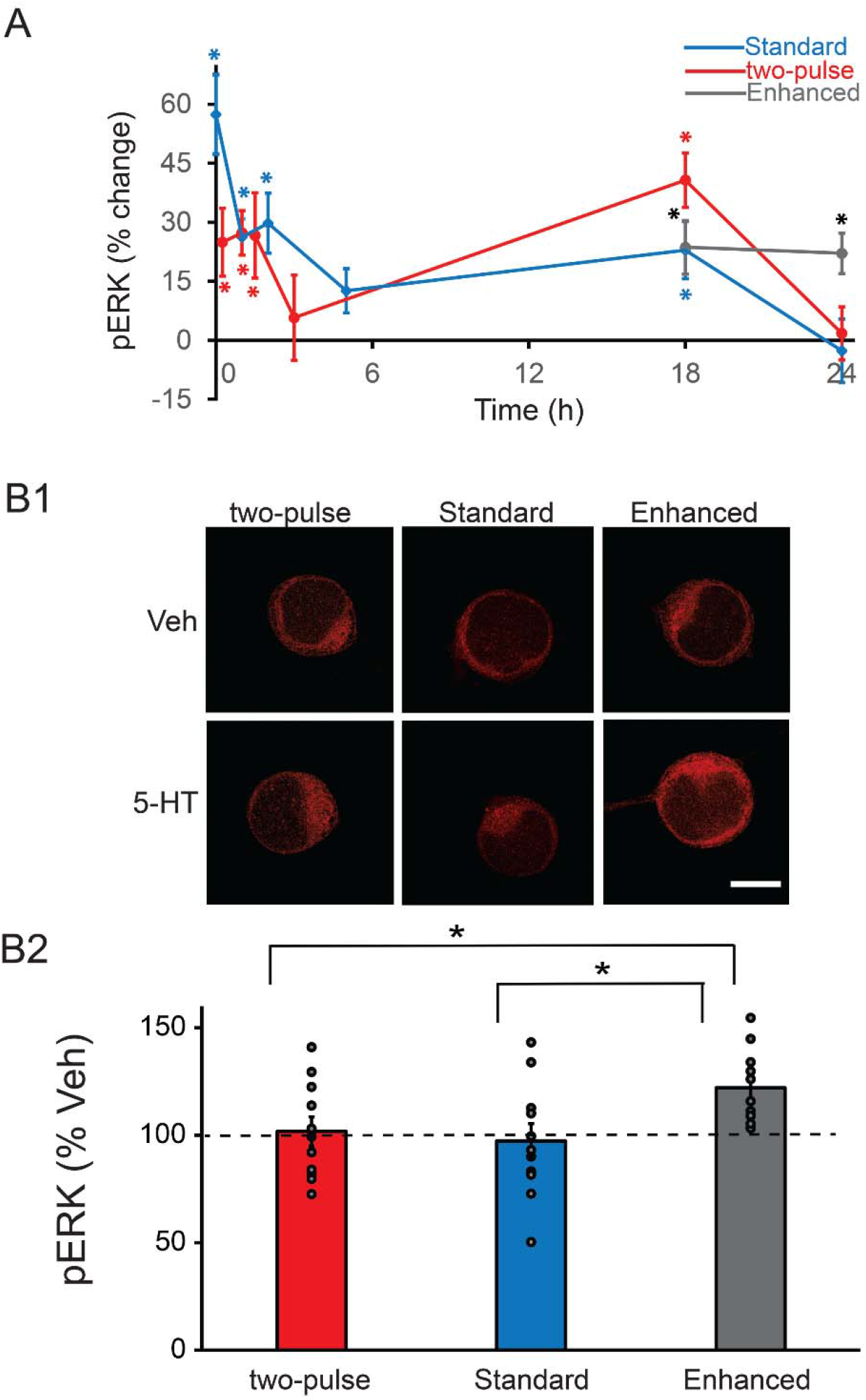
Dynamics of pERK induced by three LTF-inducing protocols. (**A**), The changes of pERK. The percent change was calculated as the change of pERK level after 5-HT compared to time-matched vehicle control. A late increase of pERK at ~18 h was induced by the two-pulse protocol (red) or Standard protocol (blue), and then followed by a return to basal level at 24 h. However, the late increase of pERK induced by the Enhanced protocol remained elevated at 24 h (black). (**B**), Changes of pERK at 24 h after different 5-HT protocols. (**B1**), Representative confocal images of pERK immunostaining in SNs. B2, Summary data. pERK induced by the Enhanced protocol was significantly greater than pERK induced by the other two protocols. Scale bar in **B1** is 40 μm. Bar heights in **B2** represent the mean, small bars represent standard error of the mean (SEM). * p< 0.05.

In the experiments to make multiple comparisons of pERK at 24 h between groups treated with three different 5-HT protocols, one-way ANOVA and the *post hoc* Student-Newman–Keuls (SNK) method were used on raw data (Fig. 2B).

In the experiments to make multiple comparisons between groups treated with 5-HT and inhibitors, repeated measures one-way (RM) ANOVA and the *post hoc* Student-Newman–Keuls (SNK) method were used on raw data (Figs. 3–4), except for data displaying a non-normal distribution (Figs. 3A3, 4B-C), which used Friedman repeated measures analysis of variance on ranks and the *post hoc* SNK method.

**Figure 3.**
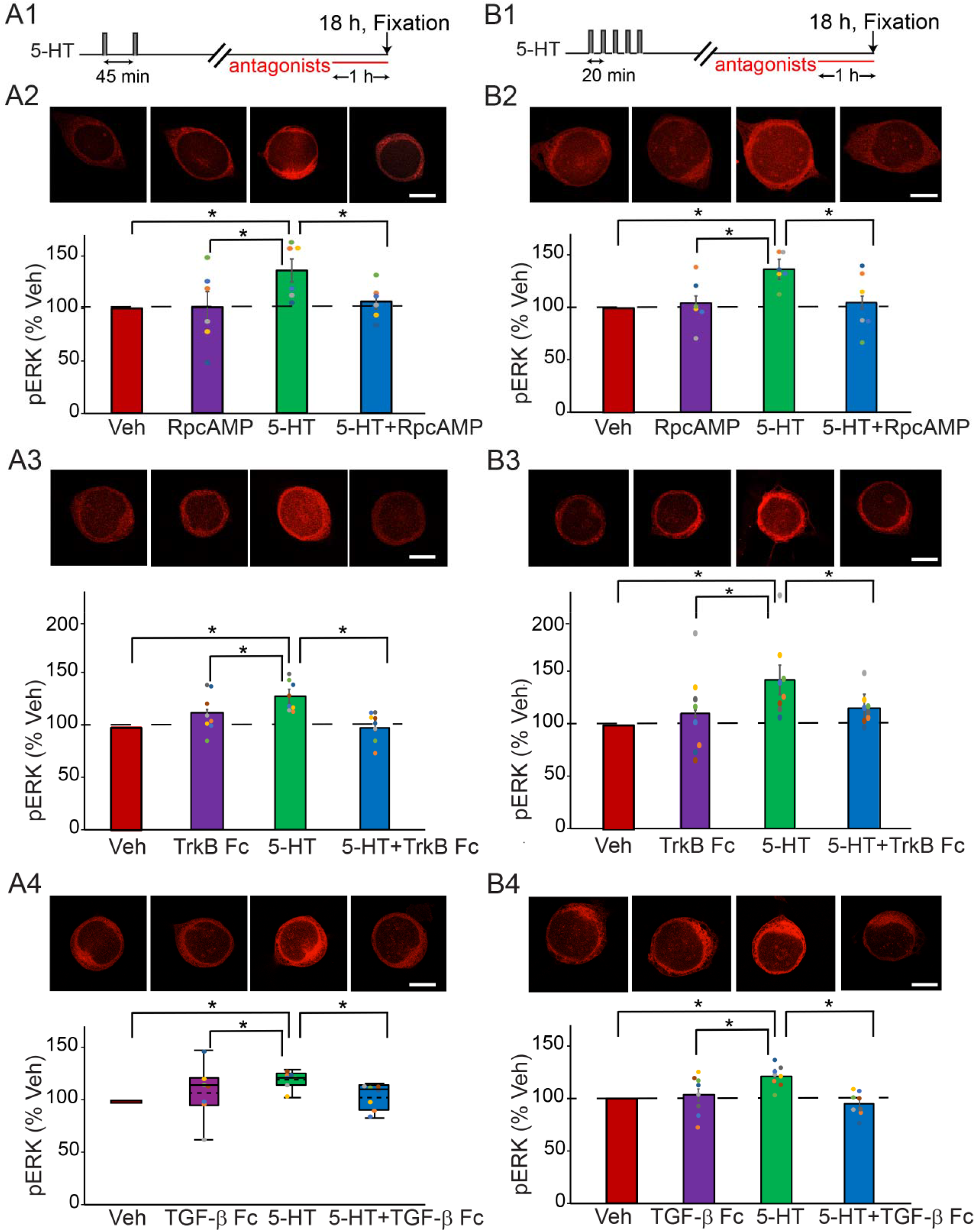
Late increase of pERK at 18 h after the two-pulse (**A**), or the Standard protocol (**B**) was dependent on PKA, TrkB, and TGF-β. (**A1**), Protocol for applying the antagonists after two-pulse protocol. (**A2**), Representative confocal images and summary data of pERK in SNs at 18 h after 2 pulses of 5-HT, in the absence or presence of PKA inhibitor RpcAMP. RpcAMP significantly decreased pERK induced by 5-HT (n =6). (**A3**), Representative confocal images and summary data of pERK at 18 h after 2 pulses of 5-HT, in the absence or presence of TrkB inhibitor TrkB Fc. TrkB Fc significantly decreased pERK induced by 5-HT (n =8). (**A4**), Representative confocal images and summary data of pERK at 18 h after 2 pulses of 5-HT, in the absence or presence of TGF-β inhibitor TGF-β RII Fc (TGF-β Fc). TGF-β RII Fc significantly decreased pERK induced by 5-HT (n =7). (**B1**), Protocol for applying the antagonists after Standard protocol. (**B2**), Representative confocal images and summary data of pERK in SNs at 18 h after 5 pulses of 5-HT, in the absence or presence of PKA inhibitor RpcAMP. RpcAMP significantly decreased pERK induced by 5-HT (n =6). (**B3**), Representative confocal images and summary data of pERK at 18 h after 5 pulses of 5-HT, in the absence or presence of TrkB inhibitor TrkB Fc. TrkB Fc significantly decreased pERK induced by 5-HT (n =8). (**B4**), Representative confocal images and summary data of pERK at 18 h after 5 pulses of 5-HT, in the absence or presence of TGF-β inhibitor TGF-β RII Fc. TGF-β RII Fc significantly decreased pERK induced by 5-HT (n =8). All scale bars are 40 μm. Bar heights in **A2-3**, **B2-B4** represent the mean, small bars represent standard error of the mean (SEM). Data in **A4** are not normal distributed, thus presented by box-and-whisker plots. The median is indicated by the solid line in the interior of the box. The mean is indicated by the dashed line in the interior of the box. The lower end of the box is the first quartile (Q1). The upper end of the box is the third quartile (Q3). The ends of the vertical lines (whiskers) are the maximum and minimum values of nonoutliers. * p< 0.05

**Figure 4.**
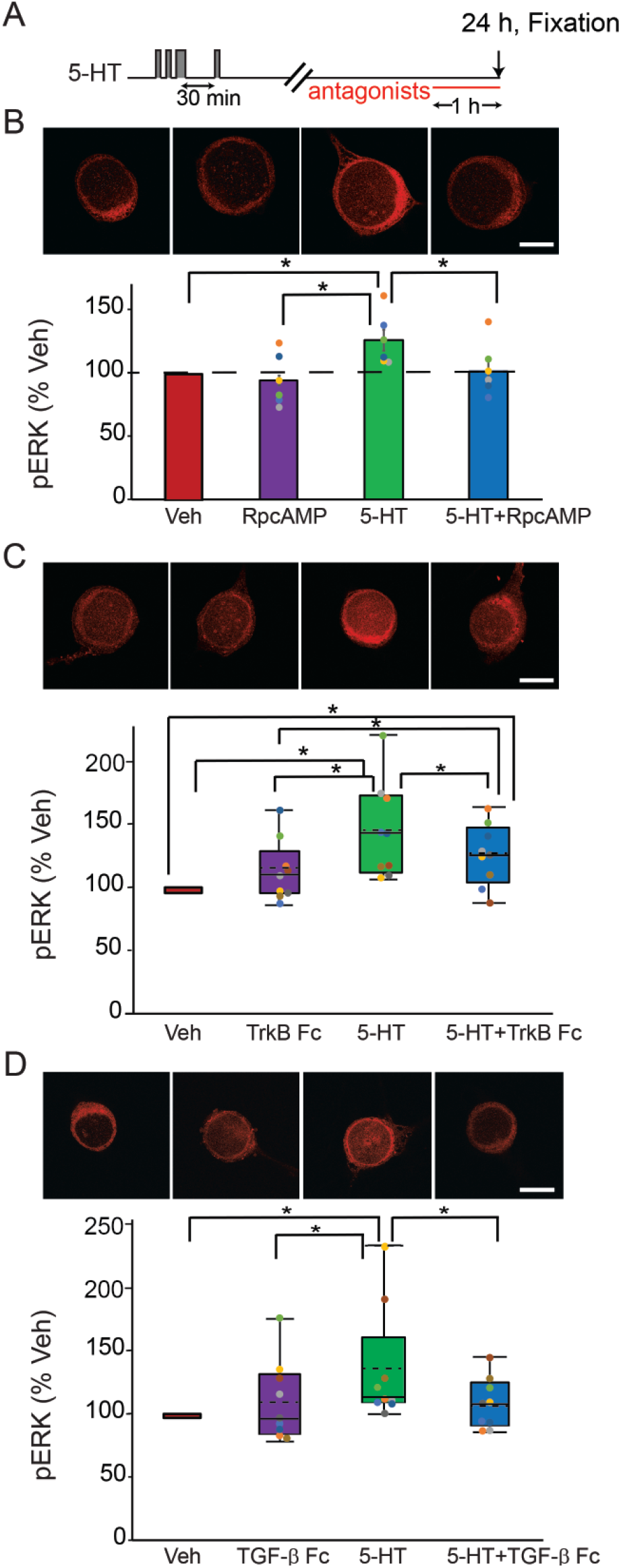
Late increase of pERK at 24 h after the Enhanced protocol was dependent on PKA and TGF-β.. (**A**), Protocol for applying the antagonists after Enhanced protocol. (**B**), Representative confocal images and summary data of pERK in SNs at 24 h after 5 pulses of 5-HT, in the absence or presence of PKA inhibitor RpcAMP. RpcAMP significantly decreased pERK induced by 5-HT (n =6). (**C**), Representative confocal images and summary data of pERK at 24 h after 5 pulses of 5-HT, in the absence or presence of TrkB inhibitor TrkB Fc. TrkB Fc significantly decreased pERK induced by 5-HT (n =9). (**D**), Representative confocal images and summary data of pERK at 24 h after 5 pulses of 5-HT, in the absence or presence of TGF-β inhibitor TGF-β RII Fc. TGF-β RII Fc significantly decreased pERK induced by 5-HT (n =9). All scale bars are 40 μm. Bar heights in **B** represent the mean, small bars represent standard error of the mean (SEM). Data in **C-D** are not normally distributed, thus presented by box-and-whisker plots. The median is indicated by the solid line in the interior of the box. The mean is indicated by the dashed line in the interior of the box. The lower end of the box is the first quartile (Q1). The upper end of the box is the third quartile (Q3). The ends of the vertical lines (whiskers) are the maximum and minimum values of nonoutliers. * p< 0.05

All of the electrophysiological and imaging experiments were conducted in a blind manner so that the investigator analyzing the data was unaware of the treatment the cells received. A p value less than 0.05 was considered to represent statistical significance. All data are available upon request.

## RESULTS

### Two-pulse 5-HT protocol induces LTF

LTM can be induced *in vivo* with only two spaced trials if the second tail shock is administered 45 min, but not 15 or 60 min, after the first (Philips et al. 2007). The first trial recruits nuclear MAPK activity that establishes a unique molecular context involving the recruitment of CREB1 kinase and CCAAT/enhancer binding protein (C/EBP), which is required for the induction of LTM (Philips et al. 2013). This LTM-inducing protocol was also able to produce LTF of the sensorimotor synapse (Fig. 1). EPSP amplitudes were measured before (pretest) and 24 h after (posttest) two, 5-min pulses of 5-HT with an ISI of 45 min (Fig. 1A). Two groups of preparations were examined, a two-pulse group and a vehicle control (Control) group, which received two 5-min applications of solution without 5-HT (45-min ISI). Sample recordings are illustrated in Fig. 1B and summary data are illustrated in Fig. 1C. The two-pulse treatment lead to a significant enhancement of the EPSPs (Veh, −2 ± 7 %, n = 7; two-pulse, 47 ± 13 %, n = 7; t_12_ = 22.06, p = 0.006).

### Two-pulse protocol and Standard protocol induce two waves of increased pERK within 24 h after 5-HT treatment

Exposure of SNs to different protocol of 5-HT treatment leads to activation of ERK (Kopec et al. 2015; Martin et al. 1997; Michael et al. 1998; Philips et al. 2013; Zhang et al. 2017, 2021), but there is a lack of comparison of detailed dynamics of long-term activation of ERK by different protocols and how ERK activity is regulated by growth factors and kinases that interact with MAPK pathways. Therefore, we used immunofluorescence to examine the dynamics of phosphorylation of ERK induced by two LTF-inducing protocols. We measured levels of pERK 15 min, 1 h, 1.5 h, 3 h, 18 h, and 24 h after two pulses of 5-HT with ISI of 45 min. The early time points within 3 h were selected based on previous studies of the two-pulse protocol (Kopec et al. 2015; Miranda et al. 2019; Shobe et al. 2016). The late time points of 18 h and 24 h were selected because previous late kinase activation might contribute to differences between the different protocols.

The two-pulse protocol induced two waves of increase in pERK (Fig. 2A, red line). ERK phosphorylation increased 15 min (24.9 ± 8.6%, n = 15), 1 h (27.3 ± 5.6%, n = 8), and 1.5 h (26.6 ± 10.8%, n = 11) after treatment. pERK returned towards basal level at 3 h (5.7 ± 10.9%, n = 10). Surprisingly, a late increase in pERK was detected at 18 h (40.7 ± 6.9%, n = 8), and then it returned to basal level at 24 h (1.8 ± 6.8%, n = 11). Statistical analyses (Wilcoxon Signed Rank Test, Bonferroni correction) revealed that the increases at 15 min, 1 h, 1.5 h, and 18 h after 5-HT were significant compared to time-matched Veh controls (15 min, z = 2.613, p = 0.042; 1 h, z = 2.521, p = 0.048; 1.5 h, z = 2.667, p = 0.03; 18 h, z = 2.521, p = 0.048), whereas the changes in pERK levels at 3 h and 24 h were both not (3 h, z = 0.153, p = 5.532 after Bonferroni corrections; 24 h, z = 0.0889, p = 5.796 after Bonferroni corrections).

Interestingly, the Standard protocol of five pulses of 5-HT with regular ISIs of 20 min induced similar pattern of dynamic changes in pERK (Fig. 2A, blue line). ERK phosphorylation increased immediately (57.4 ± 9.9%, n = 10), 1 h (26.3 ± 4.6%, n = 9), and 2 h (29.8 ± 7.6%, n = 17) after treatment. pERK returned towards basal level at 5 h (12.6 ± 5.6%, n = 10), followed by a late increase at 18 h (22.9 ± 7.3%, n = 8), and then a return to basal level at 24 h (−2.7 ± 8.1%, n = 11). Statistical analyses (Wilcoxon Signed Rank Test, Bonferroni correction) revealed that the increases immediately, 1 h, 2 h, and 18 h after 5-HT were significant compared to Veh controls (0 min, z = 2.803, p = 0.012; 1 h, z = 2.666, p = 0.024; 2 h, z = 2.627, p = 0.042; 18 h, z = 2.521, p = 0.048), whereas those at 5 h (z = 1.682, p = 0.63), and 24 h (z = −0.415, p = 4.404 after Bonferroni corrections) were not.

### Enhanced protocol leads to elevation of pERK at 24 h post-training

We hypothesized that the Enhanced protocol might produce stronger and more persistent LTF if it led to greater activation of pERK at later times. We focused on the 18 and 24-h time points. As with the Standard and two-pulse protocols, pERK was elevated at 18 h (23.6± 6.8%, n = 7) (Fig. 2A, black line). However, in contrast to both the two-pulse and Standard protocols, pERK remained elevated at 24 h (22.1 ± 5.2%, n = 11). Statistical analyses (Wilcoxon Signed Rank Test, Bonferroni correction) revealed that the increases at 18 h and 24 h after 5-HT were both significant compared to Veh controls (at 18 h, z = 2.366, p = 0.032; at 24 h, z = 2.934, p = 0.002).

To further confirm the differences in ERK activation at 24 h among three protocols, we directly compared pERK levels at 24 h following the three protocols (Fig. 2B), using the same data in Fig. 2A. A one-way ANOVA revealed a significant overall effect of the treatments (F_2,30_ = 3.804, p = 0.034). Subsequent pairwise comparisons (Student-Newman-Keuls) revealed that the Enhanced group was significantly different from the two-pulse group (q = 3.003, p = 0.042), and from the Standard group (q = 3.658, p = 0.038). No significant difference was detected between the two-pulse group and the Standard group (q = 0.655, p = 0.647).

### PKA, TrkB and TGF-β pathways mediate the late increase of pERK after the two-pulse and Standard 5-HT protocols

Previous studies suggest that the sustained increase of pERK 1 h after the two-trial protocol is, at least partially, dependent on PKA, TrkB and TGF-β pathways (Kopec et al. 2015; Zhang et al. 2021). We investigated here whether the late increase of pERK 18 h after 5-HT protocols is also dependent on these pathways.

#### PKA pathway after the two-pulse protocol

To examine the role of the PKA pathway in the late activation of ERK, the PKA inhibitor RpcAMP was applied to SNs 17 h post – 5-HT (Fig. 3A1). The cells were fixed 1 h later for immunofluorescence analysis. 5-HT led to an increase (36.2 ± 10.9%) in levels of pERK (n = 6) (Fig. 3A2). This increase was blocked (6.1 ± 6.9%) by RpcAMP (n = 6). A one-way RM ANOVA revealed a significant overall effect of the treatments (F_3,15_ = 4.273, p = 0.023, n = 6). Subsequent pairwise comparisons (Student-Newman-Keuls) revealed that the 5-HT alone group was significantly different from the Veh group (q = 4.417, p = 0.032, n = 6), from the 5-HT+RpcAMP group (q = 3.604, p = 0.022, n = 6), and from the RpcAMP alone group (q = 4.209, p = 0.024, n = 6). No significant difference was observed between RpcAMP alone and Veh (q = 0.208, p = 0.885, n = 6), between 5-HT+RpcAMP and Veh (q = 0.813, p = 0.836, n = 6), or between 5-HT+RpcAMP and RpcAMP alone (q = 0.605, p = 0.675, n = 6). Thus RpcAMP suppressed pERK elevation at 18 h after the two-pulse protocol, indicating that the late phase of activation is, at least partially, dependent on the PKA pathway.

#### TrkB pathway after the two-pulse protocol

To examine the involvement of the TrkB pathway in the late activation of ERK, TrkB Fc was applied 17 h post 5-HT and cells were fixed 1 h later for immunofluorescence. 5-HT led to an increase (23.9 ± 4.9%) in levels of pERK (n = 8) (Fig. 3A3). This increase was blocked (−3.7 ± 4.8%) in the presence of TrkB Fc (n = 8). A one-way RM ANOVA revealed a significant overall effect of the treatments (F_3,21_ = 9.501, p < 0.001, n = 8). Subsequent pairwise comparisons (Student-Newman-Keuls) revealed that the 5-HT alone group was significantly different from the Veh group (q = 5.986, p = 0.001, n = 8), from the 5-HT+TrkB Fc group (q = 6.934, p <0.001, n = 8), and from the TrkB Fc alone group (q = 3.746, p =0.015, n = 8). No significant difference was observed between TrkB Fc alone and Veh (q = 2.240, p = 0.128, n = 8), between 5-HT+TrkB Fc and Veh (q = 0.948, p = 0.51, n =8), or between 5-HT+TrkB Fc and TrkB Fc alone (q = 3.187, p = 0.085, n=8). Thus TrkB Fc suppressed pERK elevation at 18 h after the two-pulse protocol.

#### TGF-β pathway after the two-pulse protocol

To examine the role of the TGF-β pathway in the late activation of ERK, TGF-β RII Fc was applied 17 h post 5-HT and cells were fixed 1 h later for immunofluorescence. 5-HT led to an increase (19.6 ± 3.0%) in levels of pERK (n = 7) (Fig. 3A4). This increase was blocked (3.2 ± 4.5%) in the presence of TGF-β RII Fc (n = 7). A Friedman repeated measures analysis of variance on ranks revealed a significant overall effect of the treatments (chi-square= 10.714 with 3 degrees of freedom, p = 0.013, n = 7). Subsequent pairwise comparisons (Student-Newman-Keuls) revealed that the 5-HT alone group was significantly different from the Veh group (q = 4.914, p <0.05, n = 7), from the 5-HT + TGF-β RII Fc group (q = 4.099, p <0.05, n = 7), and from the TGF-β RII Fc alone group (q = 3.742, p <0.05, n = 7). No significant difference was observed between TGF-β RII Fc alone and Veh (q = 3.207, p > 0.05, n = 7), between 5-HT+ TGF-β RII Fc and Veh (q = 0.535, p > 0.05, n = 7), or between 5-HT+TGF-β RII Fc and TGF-β RII Fc alone (q = 2.646, p > 0.05, n=7). Thus TGF-β RII Fc suppressed pERK elevation at 18 h after the two-pulse protocol.

We next examined possible signaling pathways involved in the late increase of pERK 18 h after the <u>Standard</u> protocol (Fig. 3B1).

#### PKA pathway after the Standard protocol

5-HT led to an increase (34.1 ± 6.1%) in levels of pERK at 18 h (n = 6) (Fig. 3B2). This increase was blocked (2.8 ± 11.6%) in the presence of RpcAMP (n = 6). A one-way RM ANOVA revealed a significant overall effect of the treatments (F_3,15_ = 5.900, p = 0.007, n = 6). Subsequent pairwise comparisons (Student-Newman-Keuls) revealed that the 5-HT alone group was significantly different from the Veh group (q = 4.702, p = 0.012, n = 6), from the 5-HT+RpcAMP group (q = 5.196, p = 0.011, n = 6), and from the RpcAMP alone group (q = 4.591, p = 0.006, n = 6). No significant difference was observed between RpcAMP alone and Veh (q = 0.112, p = 0.938, n = 6), between 5-HT+ RpcAMP and Veh (q = 0.493, p = 0.732, n = 6), or between 5-HT+RpcAMP and RpcAMP alone (q = 0.605, p = 0.905, n=6). Thus, RpcAMP suppressed pERK elevation at 18 h after the Standard protocol.

#### TrkB pathway after the Standard protocol

5-HT led to an increase (44.3 ± 13.5%) in levels of pERK at 18 h after the Standard protocol (n = 8) (Fig. 3B3). This increase was blocked (16.8 ± 5.7%) in the presence of TrkB Fc (n = 8). A one-way RM ANOVA revealed a significant overall effect of the treatments (F_3,21_ = 8.165, p < 0.001, n = 8). Subsequent pairwise comparisons (Student-Newman-Keuls) revealed that the 5-HT alone group was significantly different from the Veh group (q = 6.612, p < 0.001, n = 8), from the 5-HT + TrkB Fc group (q = 4.110, p = 0.009, n = 8), and from the TrkB Fc alone group (q = 5.287, p = 0.003, n = 8). No significant difference was observed between TrkB Fc alone and Veh (q = 1.325, p = 0.36, n = 8), between 5-HT+TrkB Fc and Veh (q = 2.503, p = 0.204, n = 8), or between 5-HT+TrkB Fc and TrkB Fc alone (q = 1.178, p = 0.415, n = 8). Thus, TrkB Fc suppressed pERK elevation at 18 h after the Standard protocol.

#### TGF-β pathway after the Standard protocol

5-HT led to an increase (21.6 ± 3.6%) in levels of pERK at 18 h after the Standard protocol (n = 8) (Fig. 3B4). This increase was blocked (−4.7 ± 4.0%) in the presence of TGF-β RII Fc (n = 8). A one-way RM ANOVA revealed a significant overall effect of the treatments (F_3,21_ = 5.950, p = 0.004, n = 8). Subsequent pairwise comparisons (Student-Newman-Keuls) revealed that the 5-HT alone group was significantly different from the Veh group (q = 4.577, p = 0.011, n = 8), from the 5-HT+TGF-β RII Fc group (q = 5.601, p = 0.004, n = 8), and from the TGF-β RII Fc alone group (q = 3.110, p = 0.039, n = 8). No significant difference was observed between TGF-β RII Fc alone and Veh (q = 1.467, p = 0.311, n = 8), between 5-HT+ TGF-β RII Fc and Veh (q = 1.023, p = 0.478, n = 8), or between 5-HT+TGF-β RII Fc and TGF-β RII Fc alone (q = 2.491, p = 0.207, n = 8). Thus, TGF-β RII Fc suppressed pERK elevation at 18 h after the Standard protocol.

### PKA, TrkB and TGF-β pathways mediate the 24-hour increase of pERK after the Enhanced protocol

We investigated whether the sustained increase of pERK at 24 h after the Enhanced protocol is induced by the same mechanism as that underlying the increase of pERK at 18 h after the two-pulse and Standard protocols.

#### PKA pathway

To examine the role of the PKA pathway in the activation of ERK 24 h after the Enhanced protocol, RpcAMP was applied 23 h post 5-HT (Fig. 4A). SNs were fixed 1 h later for immunofluorescence. Example responses are illustrated and summary data are presented in Fig. 4B. 5-HT led to an increase (22.3 ± 8.2%) in levels of pERK (n = 6). This increase was blocked (0.1 ± 8.3%) in the presence of RpcAMP (n = 6). A one-way RM ANOVA revealed a significant overall effect of the treatments (F_3,15_ = 5.179, p = 0.012, n = 6). Subsequent pairwise comparisons (Student-Newman-Keuls) revealed that the 5-HT alone group was significantly different from the Veh group (q = 3.716, p = 0.047, n = 6), from the 5-HT+RpcAMP group (q = 3.635, p = 0.021, n = 6), and from the RpcAMP alone group (q = 5.401, p = 0.008, n = 6). No significant difference was observed between RpcAMP alone and Veh (q = 1.685, p = 0252, n = 6), between 5-HT+RpcAMP and Veh (q = 0.0809, p = 0.955, n = 6), or between 5-HT+ RpcAMP and RpcAMP alone (q = 1.766, p = 0.444, n=6). Thus RpcAMP suppressed pERK elevation at 24 h after the Enhanced protocol.

#### TrkB pathway

To examine the role of the TrkB pathway in the activation of ERK 24 h after the Enhanced protocol, TrkB Fc was applied 23 h post 5-HT. SNs were fixed 1 h later for immunofluorescence. Example responses are illustrated and summary data are presented in Fig. 4C. 5-HT led to an increase (43.6 ± 12.4%) in levels of pERK (n = 9). This increase was reduced but still significantly increased from the basal (24.6 ± 7.9%) in the presence of TrkB Fc (n = 9). Friedman repeated measures analysis of variance on ranks revealed a significant overall effect of the treatments (Chi-square= 13.533 with 3 degrees of freedom, p = 0.004, n = 9). Subsequent pairwise comparisons (Student-Newman-Keuls) revealed that the 5-HT alone group was significantly different from Veh (q = 4.648, p < 0.05, n = 9), from 5-HT+TrkB Fc (q = 3.771, p < 0.05, n = 9), and from TrkB Fc alone (q = 5.333, p < 0.05, n = 9). The 5-HT+TrkB Fc group was significantly different from Veh (q = 3.333, p < 0.05, n = 9), from TrkB Fc alone (q = 3.771, p < 0.05, n = 9). No significant difference was observed between TrkB Fc alone and Veh group (q = 0.943, p < 0.05, n = 9). These results indicate that TrkB Fc can reduce, but cannot completely block, pERK elevation at 24 h after the Enhanced protocol.

#### TGF-β pathway

To examine the role of the TGF-β pathway in the activation of ERK 24 h after the Enhanced protocol, TGF-β RII Fc was applied 23 h post 5-HT. SNs were fixed 1 h later for immunofluorescence. Example responses are illustrated and summary data are presented in Fig. 4D. 5-HT led to an increase (34.7 ± 15.2%) in levels of pERK (n = 9). This increase was blocked (7.8 ± 6.7%) in the presence of TGF-β RII Fc (n = 9). A Friedman repeated measures analysis of variance on ranks revealed a significant overall effect of the treatments (chi-square= 10.200 with 3 degrees of freedom, p = 0.013, n = 9). Subsequent pairwise comparisons (Student-Newman-Keuls) revealed that the 5-HT alone group was significantly different from the Veh group (q = 5.000, p <0.05, n = 9), from the 5-HT+TGF-β RII Fc group (q = 5.657, p <0.05, n = 9), and from the TGF-β RII Fc alone group (q = 3.873, p <0.05, n = 9). No significant difference was observed between TGF-β RII Fc alone and Veh (q = 0, p > 0.05, n = 9), between 5-HT+TGF-β RII Fc and Veh (q = 1.414, p > 0.05, n = 9), or between 5-HT+TGF-β RII Fc and TGF-β RII Fc alone (q = 1.000, p > 0.05, n = 9). Thus, TGF-β RII Fc suppressed pERK elevation at 24 h after the Enhanced protocol.

## Discussion

Immunofluorescence analysis was used to characterize the dynamics of activation of ERK up to 24 h after different 5-HT protocols, in the absence vs. presence of inhibitors. Results suggest that multiple pathways and feedback loops contributes to the long-term activation of pERK (Fig. 5). Differential pERK dynamics observed during the late consolidation phase after these protocols provided insights into the mechanism underlying the advantage of the Enhanced protocol in producing prolonged LTF.

**Figure 5.**
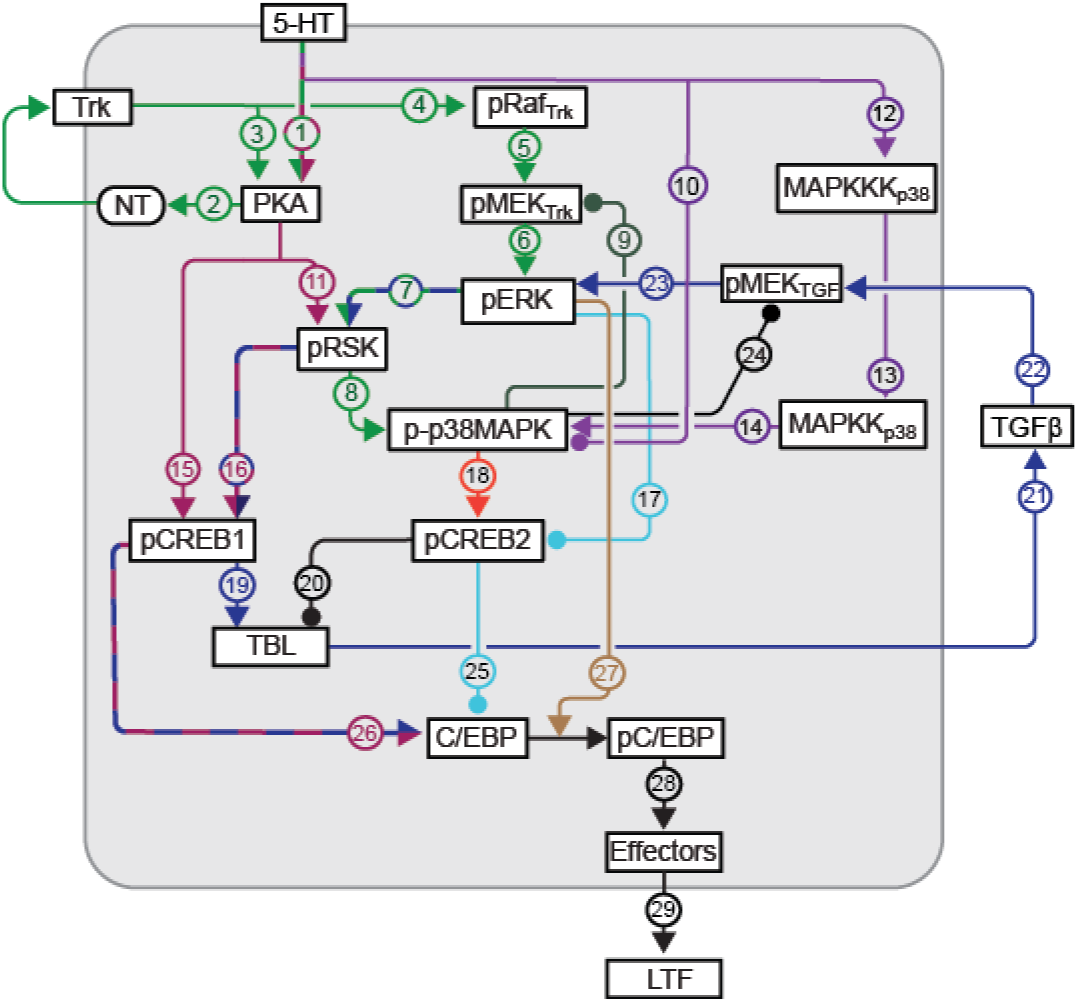
Molecular pathways in SNs that regulate LTF. Release of 5-HT initiates cascades that activate PKA, ERK, p38 MAPK and RSK. These kinase cascades interact with growth factors NT and TGF-β to induce multiple feedback loops for the sustained increase of kinase activity up to 24 h after 5-HT. Active kinases, in turn activate genes (e.g. creb1/2, c/ebp) essential for the induction of LTF, through interlocking positive and negative feedback loops that determine CREB1 and CREB2 activity. Arrowheads indicate activation, circular ends indicate repression.

### pERK has complex dynamics

Empirical studies identified three different 5-HT protocols that can induce LTF: the Standard protocol; the Enhanced protocol; and the 2-pulse protocol. The Standard and two-pulse protocols led to an immediate increase in pERK, which decayed within 5 h post treatment. A second wave of increased pERK was detected at 18 h post treatment. This late phase was blocked by RpcAMP, and by antagonists TrkB Fc and TGF-β RII Fc. PKA activity is increased at 20 h after the Standard protocol (Müller and Carew 1998). The results here suggest that PKA activity might be already elevated at 18 h, to directly or indirectly activate TrkB and TGF-β, leading to the increase of pERK (Fig. 5, pathways 1 (green/red)->2 (green)->4 (green)->5 (green)->6 (green); 1 (green/red) −>15 (red)->19 (blue)->21 (blue)->22 (blue)->23 (blue)). It is also highly likely that rich dynamics of ERK will be shared by other kinases and transcription factors. For example, complex dynamics of CREB1 and CREB2 after Standard protocol have been reported in previous study (Liu et al. 2008, 2011). pERK might affect the dynamics of CREB1 via RSK (Fig. 5, pathways 7 (green/blue)->16 (blue/red)), and affect the dynamics of CREB2, directly or via p38 MAPK (Fig. 5, pathways 17 (light blue), 7 (green/blue)->8 (green)->18 (orange)).

### Multiple feedback loops of the ERK, PKA, TrkB and TGF-β contribute to the dynamics of long-term ERK activation

The model of Fig. 5 illustrates aspects of the complex network of pathways that induces LTF. One pulse of 5-HT activates PKA, which increases the presynaptic release of NT (green pathways 1 (green/red)->2 (green)). NT binds its receptor TrkB, resulting in a slow activation of the Raf-MEK-ERK pathway (green pathways 4->5->6). After two or more pulses of 5-HT, TBL – TGF-β signaling is activated, likely via the transcriptional activator CREB1 which is phosphorylated by PKA (pathways 15 (red)->19 (blue)->21 (blue)) (Chin et al. 1999; 2006; Ohta et al. 2008; Zhang et al. 1997, 2011). TGF-β in turn enhances the activation of ERK and CREB1, forming a positive feedback loop (pathways 19 (blue)->21 (blue)->22 (blue)->23 (blue)->7 (blue/green)->16 (blue/red)->19 (blue)) (Chin et al. 2006; Kopec et al. 2015; Liu et al. 1997; Miranda et al. 2019; Zhang et al. 1997). Zhang et al. (2021) also suggests that although the TGF-β – ERK dependent feedback loop is the major contributor to sustained increase of ERK activity, transient activation of NT – TrkB – ERK dependent pathways is necessary to activate this feedback loop.

Müller and Carew (1998) observed three phases of PKA activation after 5-HT – early, intermediate, and late (lasting respectively ~5 min, ~3 h, and up to ~20 h). The intermediate phase might be due to the PKA – NT – TrkB – PKA positive feedback loop (Fig. 5, green pathways 2->3) (Jin et al. 2019). The late phase might be induced by a CREB – Ap-Uch – PKA feedback loop (Chain et al. 1999; Hegde et al. 1997). The long-term dynamics of ERK activation are unclear. In this study, we found a complex dynamic pattern of ERK activation. Two waves of increase in pERK were induced by the two-pulse and Standard protocols. The first wave returned to the basal level within 5 h after 5-HT, whereas the second wave occurred around 18 h after 5-HT. These dynamics were similar to those of PKA activation after the Standard protocol (Müller and Carew 1998). Application of inhibitors suggests that the late increases of pERK induced by the two-pulse and Standard protocols are dependent on the PKA, TrkB, and TGF-β pathways. We posit that the late phase of PKA activation directly activates the PKA – NT – TrkB – PKA positive feedback loop (Fig. 5, green pathways 2->3), and indirectly activates the TGF-β – ERK feedback loop via phosphorylation of CREB1 (Fig. 5, pathways 15 (red)->19 (blue)->21 (blue)- >22 (blue)). Both feedback loops contribute to the late increase of pERK at 18 h (Fig. 5, green pathways 4>5->6; blue pathways 22->23). Sustained increase of ERK and PKA pathways converge to regulate CREB1 and CREB2 (pathways 15 (red), 7 (green/blue)->16 (red/blue), 7 (green/blue)->8 (green)->18 (orange), 17 (light blue)) (Bartsch et al. 1995, 1998), which subsequently increase the expression of C/EBP, essential for the induction of LTF (pathways 25 (light blue), 26 (red/blue)) (Alberini et al. 1994; Alberini and Kandel 2014; Guan et al. 2002; Mohamed et al. 2005).

### Putative mechanism underlying the superiority of the Enhanced protocol in producing prolonged LTF

We found that pERK remained elevated at 24 h after the Enhanced protocol, whereas it returned to basal 24 h after the two-pulse and Standard protocols. In Fig. 5, three feedback loops directly or indirectly contribute to the increase of pERK after 18 h: PKA-NT-TrkB feedback loop, PKA-CREB1 feedback loop (via ApUch), and ERK - CREB1 - TGF-β feedback loop. pERK returned to basal 24 h after the two-pulse and Standard protocols might be due to the effect of ERK-RSK-p38 MAPK negative feedback loop negative feedback loop (Zhang et al. 2021) (Fig. 5, pathways 7 (green/blue)->8 (green)->9 (dark green)->6 (green)). Our experiments suggest the pERK increase at 24 h can be blocked by inhibiting PKA or TGF-β, but not completely blocked by inhibiting TrkB. Thus, the TGF-β – ERK feedback loop appears to play a greater role in the increase of pERK at 24 h. The Enhanced protocol might induce a stronger TGF-β – ERK feedback loop than do the two-pulse and Standard protocols, resulting in ERK activation prolonged at least up to 24 h, resistant to ERK-p38 MAPK negative feedback. This hypothesis is consistent with our previous finding that the Enhanced protocol induces greater CREB1 activity at 18 h than the Standard protocol (Zhang et al. 2011). Stronger CREB1 activity at 18 h might lead to a sustained activation of the TGF-β – ERK feedback loop at 24 h. Stronger CREB1 will enhance the expression of C/EBP (Fig. 5, pathway 26 (red/blue)), and stronger pERK will enhance the activity of C/EBP (Fig. 5, brown pathway 27), which will lead to an enhanced and prolonged LTF (Fig. 5, black pathways 28->29).

### Late wave of increase in ERK might provide an optimal time to apply a second block of 5-HT

LTF was originally defined as synaptic facilitation occurring 24 h after training (Montarolo et al. 1986). It has now been measured at 48 h, 72 h, and even 7 d after training (Casadio et al. 1999; Hu et al. 2011; Liu et al. 2011; Miniaci et al. 2008). Two or more blocks of training, or of 5-HT treatment, have been found to prolong LTM and LTF. Each block consists of a Standard protocol and the interval between blocks is ~24 h. This protocol produces long lasting LTM or LTF as assessed 7 days later (Hu et al. 2011, 2015). These data suggest that some “trace” of kinase activity remains elevated, such that the trace from the first block of trials enhances the effects of the second block. In these studies as well as others, the second block has been given at a fixed time (24 h) after the first. Here, we found that ERK was activated ~18 h after all three LTF-inducing protocols, but remained activated at 24 h only after the Enhanced protocol. These results support the existence of a “trace” of active kinase after the first block of trials that might facilitate the effects of a second block of trials.

Because many of the molecular processes involved in LTF and late LTP are conserved between *Aplysia* and mammals, the findings of this study highlight the importance of examining the dynamics of kinase cascades in neurons involved in long-term memory in vertebrates. A knowledge of these dynamics may provide insights into mechanisms underlying late LTP, and help design protocols to enhance memory in mammals.

## ACKNOWLEDGEMENTS

The authors thank E. Kartikaningrum and J. Zhong for preparing cultures, and C. Neveu for statistical advice and help with the illustrations. This study was supported by NIH grant NS019895.

